# A posterior approach to correct for focal plane offsets in lattice light sheet structured illumination microscopy

**DOI:** 10.1101/2024.04.26.590138

**Authors:** Yu Shi, Tim A. Daugird, Wesley R. Legant

## Abstract

**Significance:** Lattice light sheet structured illumination microscopy (latticeSIM) has proven highly effective in producing 3D images with super resolution rapidly and with minimal photobleaching. However, due to the use of two separate objectives, sample-induced aberrations can result in an offset between the planes of excitation and detection, causing artifacts in the reconstructed images.

**Aim:** We introduce a posterior approach to detect and correct for the axial offset between the excitation and detection focal planes in latticeSIM and provide a method to minimize artifacts in the reconstructed images.

**Approach:** We utilized the residual phase information within the overlap regions of the laterally shifted structured illumination microscopy (SIM) information components in frequency space to retrieve the axial offset between the excitation and the detection focal planes in latticeSIM.

**Results:** We validated our technique through simulations and experiments, encompassing a range of samples from fluorescent beads to subcellular structures of adherent cells. We also show utilizing transfer functions with the same axial offset as that which was present during the data acquisition results in reconstructed images with minimal artifacts and salvages otherwise unusable data.

**Conclusion:** We envision that our method will be a valuable addition to restore image quality in latticeSIM datasets even for those acquired under non-ideal experimental conditions.

## 1. Introduction

Lattice light sheet microscopy (LLSM) has been widely applied in biological imaging across various scales, spanning from biomolecules to embryos^1–3^. This technique offers several advantages over epifluorescence or confocal microscopy, including minimal out-of-focus fluorescence, reduced photobleaching and phototoxicity, and enhanced imaging speed. By utilizing the interference pattern from multiple beams, LLSM improves beam uniformity and axial resolution compared to Gaussian beams^4,5^. Typically, to ensure uniform sample illumination and maximize imaging speed, lattice light sheets are laterally dithered to average out modulations due to the interfering beams. However, by stepping the lattice in discrete increments rather than dithering, the same light sheet can be utilized for super resolution SIM yielding improved lateral and axial resolution^2^. LatticeSIM also more thoroughly fills out the optical transfer function (OTF) axially compared with dithered lattice illumination which results in better image quality. However, due to the fixed objective orientation and the lower numerical aperture (NA) of the excitation objective compared to the detection objective, latticeSIM has less resolution improvement than 3D SIM and these improvements are restricted to only a single orientation. Nevertheless the lower photobleaching and phototoxicity of latticeSIM has proven useful for imaging a variety of biological samples^2,6^.

In addition to the limitations described above, the dual-objective requirement for latticeSIM also introduces the risk of potential misalignment between the excitation pattern and the detection focal plane. Although this is common to all dual-objective light sheet systems, focus mismatch is particularly detrimental for latticeSIM due to the high NA of excitation and detection and the complexities of the classical SIM reconstruction algorithm^7^ which can lead to artifacts in the final image^8^.

Although there are a variety of autofocusing routines^9–12^ to correct mismatch in axial focus between excitation and detection objectives, these methods face three primary limitations. First, autofocusing often necessitates a separate imaging routine that must be interspersed with a normal timelapse scan. This leads to additional photobleaching and photoxicity and slower acquisition rates. Second, sample-induced axial misalignment might vary across the biological sample and even within a single field of view, therefore requiring multiple iterations of autofocusing in different regions. And finally, intensity-based optimizations may not work correctly for the periodic illuminations used in latticeSIM. Hence, there is motivation to develop a posterior approach to detect and correct for misalignments caused by system drift or sample-induced aberrations in latticeSIM. If successful, such an approach could salvage misaligned data, adapt to changes in live samples on the fly, and avoid the need for additional imaging routines.

Inspired by previous publications in opposed objective, interferometric structured illumination^13,14^, we propose a method to ascertain the axial mismatch between excitation and detection focal planes in latticeSIM imaging purely from the raw datasets. By measuring the residual phase within the overlap regions among different laterally shifted frequency components, we establish that it is possible to posteriorly determine the axial offset between the illumination pattern and the detection focal plane. We demonstrate the efficacy of this method through simulations and validation using fluorescent beads and biological samples. Once the offset has been determined, we show that reconstructing with transfer functions acquired with the same axial offset as retrieved by our method successfully mitigates artifacts arising from misalignment in the raw data.

## 2. Methods

### 2.1 Theory

For clarity, we adopt here a similar notation to that used in Gustafsson et al^7^. In fluorescence microscopy, the observed raw data *D*(***r***) is the convolution of the sample-emitted fluorescence *E*(***r***) and microscope’s detection point spread function *H*(***r***).

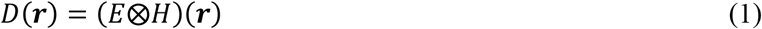

The emitted fluorescence is the product of the fluorescently labelled sample structure S(r) and the excitation point spread function *I*(***r***).

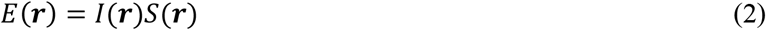

This can also be written in frequency space as 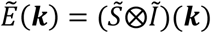, where 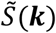 and *Ĩ*(***k***) are the Fourier transforms of S(**r**) and I(**r**).

The latticeSIM excitation pattern I(**r**) is formed by the interference of multiple beams at the back pupil of the excitation objective. In the case of the hexagonal lattice pattern, the six beams at the back pupil (**Fig 1a and b**) will interfere and form an excitation pattern at the sample plane that is the squared magnitude of the Fourier transform of the pupil function (**Fig 1c**). Similarly, the excitation optical transfer function *Ĩ* (***k***) is the Fourier transform of I(**r**) or the autocorrelation function of the pupil function (**Fig 1d**). As is the case for conventional 3D SIM, the excitation pattern can be expressed as a finite sum of “m” components, each of which can be separable into functions that depend solely on the lateral or axial coordinates in real space or frequency space respectively.

**Figure 1:**
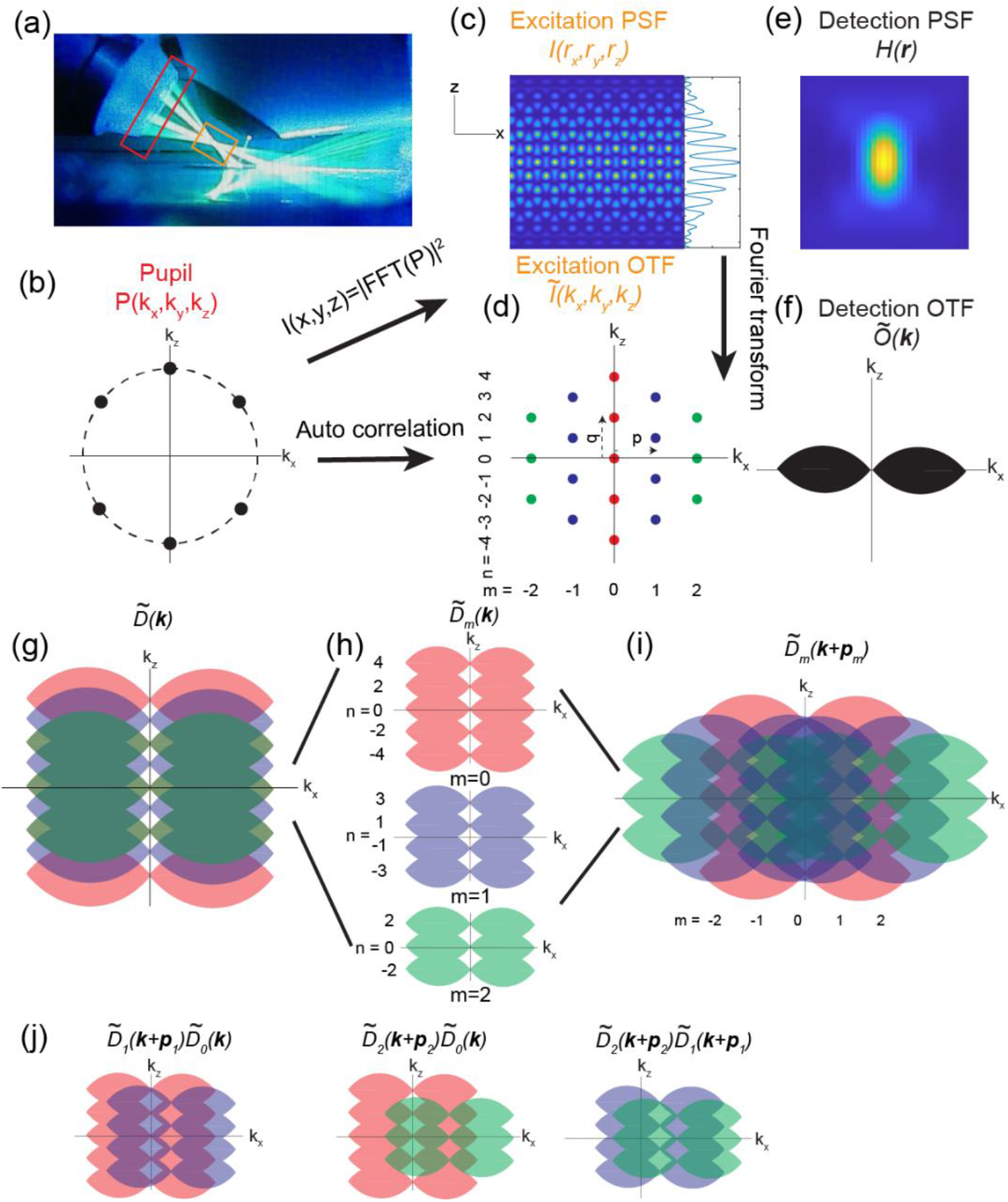
Overview of latticeSIM imaging. **(a)** Image of the latticeSIM setup. Six beams are emitted by the excitation objective (left) and form a hexagonal interference pattern. The detection objective (right) collects the emitted fluorescence. The red and orange box highlight the positions of pupil and sample planes respectively **(b)** The excitation pattern at the back pupil plane of the excitation objective. **(c)** The excitation point spread function (PSF) of the hexagonal pattern that illuminates the sample. It is the squared amplitude of the Fourier transform of the pupil function. Inset shows x-averaged intensity profile along the z axis. **(d)** The excitation optical transfer function (OTF) of the hexagonal pattern. It is a Fourier transform of the excitation PSF and an auto correlation of the pupil function. Different lateral orders (m) are coded by color. Different axial orders (n) are listed as well. **(e)** Detection PSF *H*(***r***). **(f)** Widefield detection OTF *Õ*(***k***). It is a Fourier transform of the detection PSF. G-I) Structure illumination microscopy (SIM) imaging and reconstruction process. In the SIM images 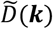 **(g)**, information components for different lateral orders are shifted to be centered at k_x_ = 0. In SIM reconstruction, these components are separated **(h)** and then shifted back to their correct position in frequency space **(i). (j)** Schematic of the overlap regions between different orders in 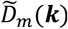, left: m = 0 and m =1; middle: m = 0 and m = 2; right: m = 1 and m = 2.

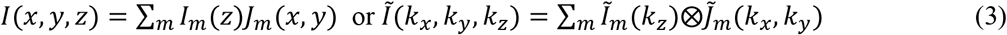

For hexagonal lattice illumination, the excitation OTF has signal at 5 lateral frequencies and m ranges from −2 to +2 as shown in **Fig 1d**.

Rewriting equation (1) with equation (2) and the separated components in the excitation PSF (3), it becomes:

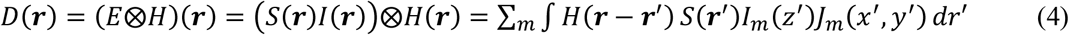

In the equation above with a spatially invariant PSF, **r’** stands for the reference frame of the specimen and **r-r’** stands for the reference frame of the objective lenses. When operating the microscope, the excitation pattern is kept fixed relative to the detection objective focal plane while the sample is translated along the “z” direction to acquire a 3D volume. Therefore, the axial components of the excitation pattern *I*_*m*_ follow the same reference frame as the objectives (same as *H*(***r*** − ***r***’)) rather than the sample, and *I*_*m*_(*z′*) can be replaced by *I*_*m*_(*z* − *z′*). We refer the reader to a full discussion in Gustafsson et al^7^ for more information about the coordinate reference frames. Equation (4) can then be written as:

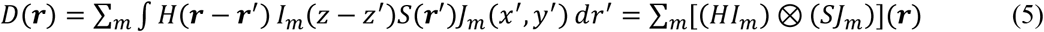

And its Fourier transform 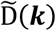 is

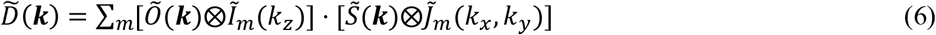

Where *Õ* (***k***) is the Fourier transform of *H*(***r***) (**Fig 1e and f**), which is the widefield OTF of the detection objective. For SIM illumination, *J*_*m*_(*x, y*) typically follows a simple harmonic where 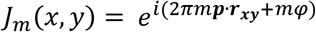 and 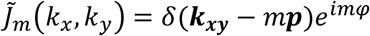. Here, *p* is the fundamental lateral frequency of the illumination pattern, *mp* are the m^th^ order harmonics, and *e*^*imφ*^ defines the lateral phase for J.

Therefore equation (6) becomes:

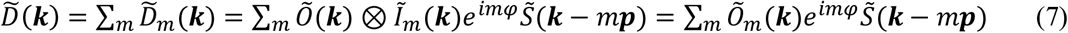

Equations 6 and 7 indicate that the resolution is improved by two methods. First, resolution is increased axially through *Õ*_*m*_(***k***) = *Õ* (***k***) ⊗ *Ĩ*_*m*_(***k***). The overall OTF becomes a series of transfer functions that are the convolution between the widefield detection OTF and the axial part of each excitation OTF order m. And second, sample information components 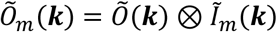 have been shifted laterally into the detection envelope by the vector p_m_ (**Fig 1g**). Note that because the high-resolution axial components are already in their correct locations in frequency space (because the sample is scanned axially as described above), the SIM reconstruction process will need to separate the different “m” components of 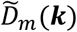 (**Fig 1h**) and then shift the high frequency components laterally back to their original positions in frequency space (**Fig 1i**), thus yielding improved resolution (see Gustafsson et al^7^ for details).

In conventional SIM imaging, the fundamental frequency ***p*** and the lateral starting phase *φ* may be different from when the transfer functions, *Õ*_*m*_(***k***) are measured and when the sample is imaged. Thus, these parameters are typically fit after data acquisition and during the reconstruction process. This is done by examining the overlap regions between different m components of 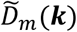 in frequency space.

More specifically, once the separated information components of the data have been isolated and shifted to their correct locations in frequency space, the frequency information of the sample 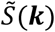 should be identical in the overlapping regions. However, as noted in eqn (7) the sample information 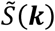 in the observed data terms 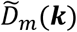 has also been scaled and phase shifted by a corresponding, order-specific transfer function *Õ*_*m*_(***k***) as a result of the physical observation process. Thus, the shifted information components 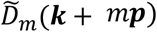 are not directly comparable to each other. In order to compensate for this, we multiply the observed data components 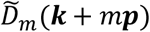 by the corresponding transfer function of the other information component in the overlap region. For example, in the overlap regions between m = 0 and m = 1, we compare

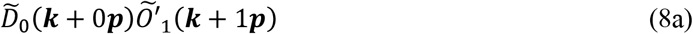

And

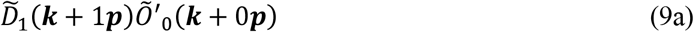

as shown in (**Fig 1j**). Here *Õ′*_*m*_(***k***) are the transfer functions obtained when measuring calibration beads at the start of an experiment, e.g. *Õ′*_*m*_(***k***) is equivalent to 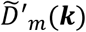 with 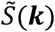 replaced by a delta function. Typically for the calibration images, the lateral phase of the excitation pattern, which depends only on the relative position of the bead used for measurement, is set to zero. 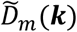 is the sample image in frequency space, and because each 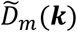 already contains a copy of *Õ*_*m*_(***k***) as described above, eqns (8a) and (9a) can each be expanded as

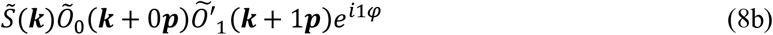

And

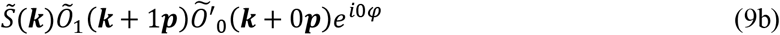

If we first assume that *Õ′*_*m*_(***k***) = *Õ*_*m*_(***k***), which means that the transfer functions when taking the sample image were identical to those obtained when taking the calibration images, then at the correct shift vector p, eqns (8) and (9) above will be identical in theory except for a constant phase offset *φ*. In practice and in the presence of noise, ***p*** is determined as the value that maximizes the cross correlation within the overlapping region. At the correct value for ***p***, the starting phase *φ* associated with 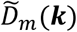 can be determined by examining the ratio between eqns (8) and (9):

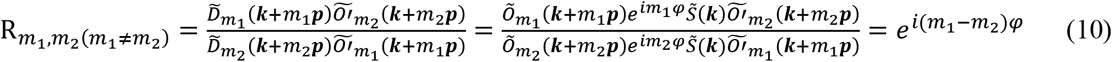

Note this ratio is independent of sample information and is constant throughout the entire overlap region. Therefore, *φ* can be extracted using complex linear regression upon the two terms, and the slope carries the information of the starting phase *φ*.

In contrast to conventional single-objective SIM, in latticeSIM the axial components in *Ĩ*_*m*_(*k*) may also acquire a phase term that is dependent on δ*z* – the physical offset between the detection and excitation reference planes. In this case, assuming that the illumination pattern along distribution, the axial components of the excitation pattern can be defined as:

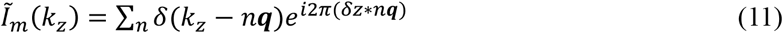

Here **q** is the fundamental frequency along the axial direction and n is a function of m. Specifically, as shown in **Fig 1d**

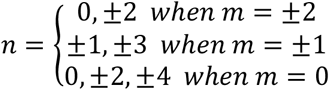

2π(δ*z* ∗ *n****q***) is the phase shift in the excitation OTF that is caused by the physical misalignment δ*z* between the excitation pattern and the detection objective focal plane (**Fig 2a and b)**. Because the overall OTF is the convolution of the detection OTF with the excitation OTF (**Fig 1g**), any phase term in the excitation OTF manifests in the overall transfer functions of the instrument (**Fig 2c**). Therefore, the amount of axial misalignment of the excitation pattern can, in theory, be extracted if we could determine the relative contribution to the phase information in 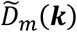 that was due to the transfer functions *Õ*_*m*_(***k***) when the image was acquired.

**Figure 2:**
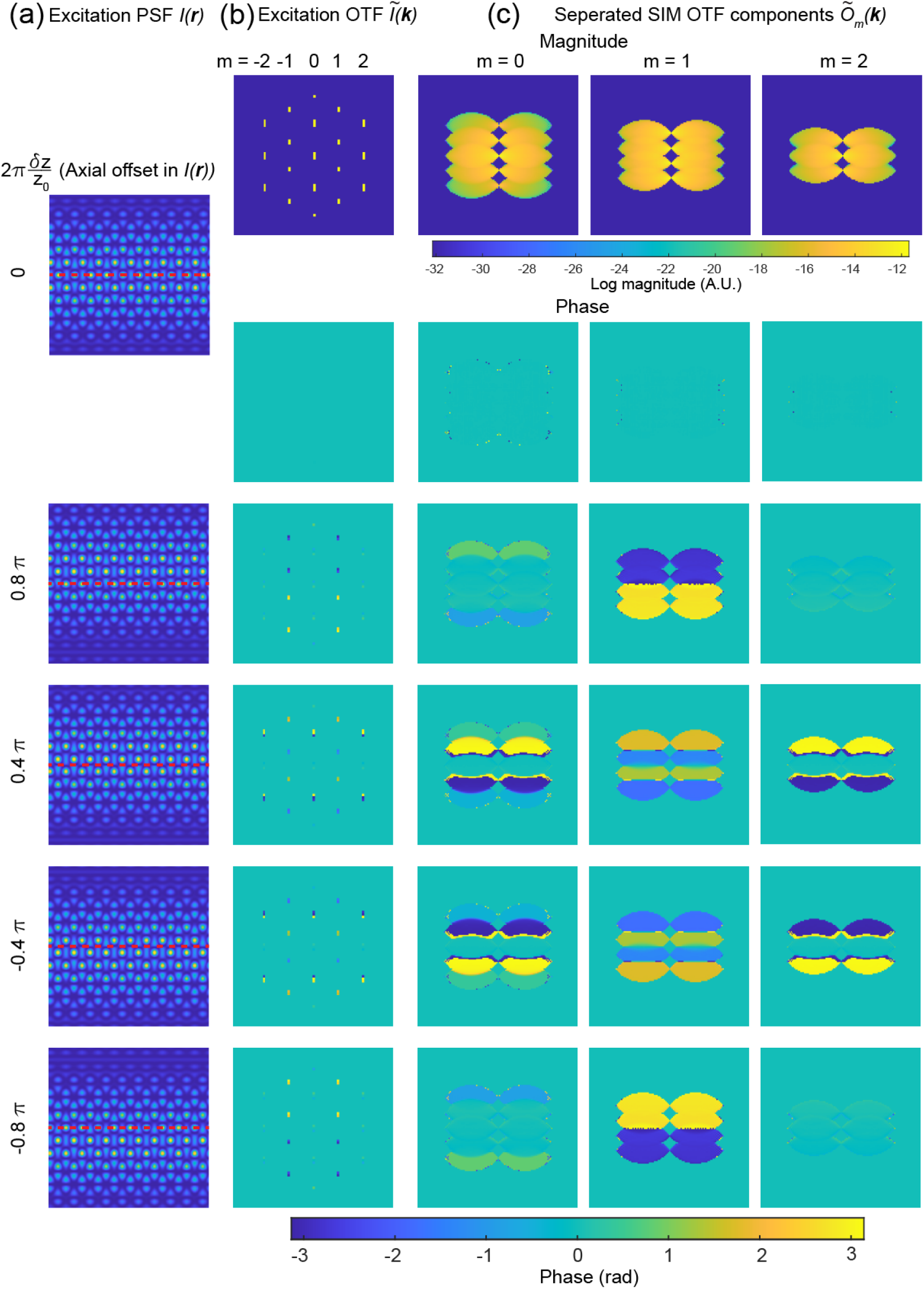
Excitation and SIM OTFs with different axial offsets. **(a)** Excitation PSF *I*(***r***) in real space. The number at the left indicates the offset from the simulated detection objective focal plane. **(b)** Excitation OTF *Ĩ* (***k***). in frequency space. The first row shows the magnitude. The rows underneath show the phase under different axial offsets. **(c)** Separated SIM transfer functions *Õ*_*m*_(***k***) (same as Fig 1h) showing the magnitude and phase under different excitation pattern axial offsets.

However, the transfer functions *Õ*_*m*_(***k***) are not typically accessible in raw images, and as with the lateral starting phase above, the axial offset may vary from when the transfer functions are calibrated at the beginning of an experiment and when a given image is acquired. For example, as shown in eqn (7), the phase of the raw images 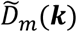 composed of eight beads (right most column in **Fig 3a**) is a mixture of illumination and sample information (**Fig 3a and b**). Therefore, to retrieve the axial misalignment, we need to first cancel out the sample information 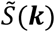 from the observed image 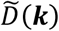. This can be achieved by the same approach described in eqn (10). If we include an axial offset which leads to phase ramp in *Ĩ*_*m*_(*k*_*z*_) as described in eqn (11), and assume that there may be a different axial offset δ*z* and δ*z*’ when imaging the sample or calibration beads respectively, then eqn (10) then becomes

**Figure 3:**
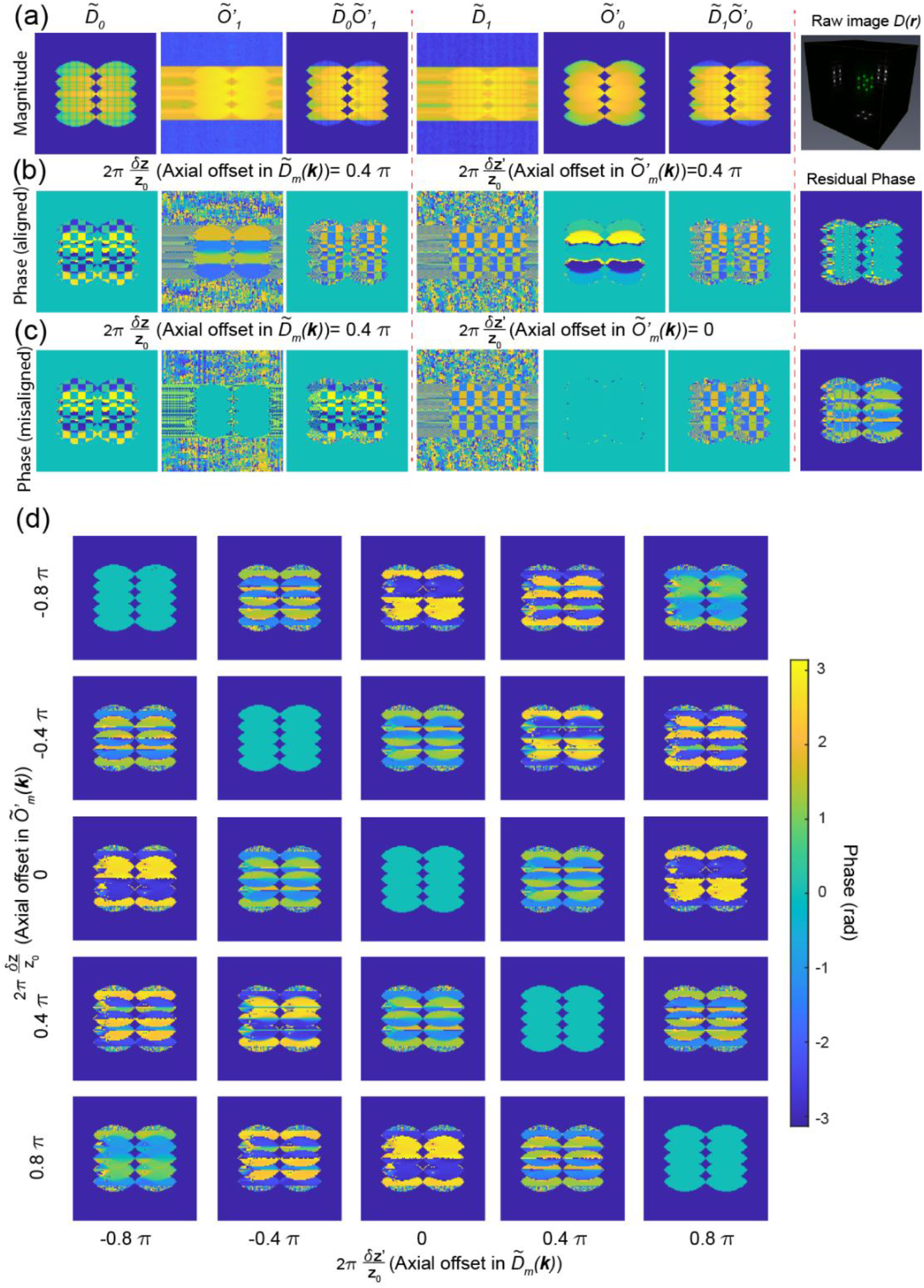
Residual phase in OTF overlaps reflects the axial offset. **(a)** Columns 1-3: Magnitude of the zeroth information component 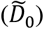, the first SIM transfer function (*Õ′*_1_) after being shifted to the correct position in frequency space, and the magnitude of the product of the two. Here, the SIM image is composed of eight beads. Columns 4-6: Magnitude of the first information component in SIM image after being shifted to the correct position in frequency space 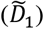, the zeroth SIM transfer function (*Õ′*_0_), and the magnitude of the product of the two. **(b)** The phase of the corresponding columns in (a). In this case, both the SIM image 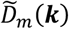 and transfer functions (*Õ′*_*m*_ (***k***)), have been simulated with a 0.4 π axial offset between the excitation and detections. The rightmost column shows the residual phase as measured via the ratio in eqn (12). **(c)** Same as **(b)** but for in this case, the transfer function *Õ′*_*m*_(***k***), does not have the corresponding 0.4 π axial offset as 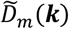. **(d)** Residual phase as computed via eqn (12) for different axial offsets between the excitation and detection planes in simulated SIM images (different columns) and simulated transfer functions (different rows).

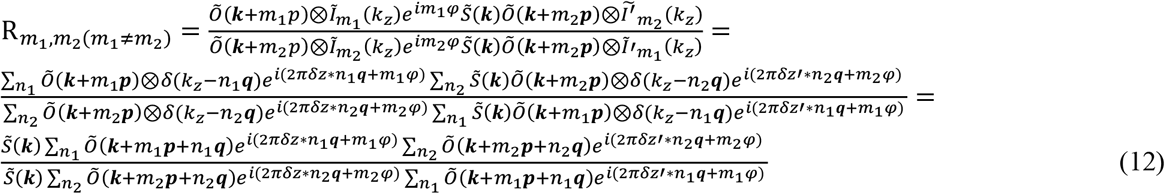

Note that here, we have expanded *Õ*_*m*_(***k***) = *Õ* (***k***) ⊗ *Ĩ*_*m*_(*k*_*z*_). We also use *Ĩ*_*m*_(*k*_*z*_) and *Ĩ′*_*m*_(*k*_*z*_) to denote the axial component of illumination when imaging the sample and the calibration beads respectively. Here we assume there is no phase contributed from the widefield detection OTF, as it can be corrected by proper alignment or adaptive optics. Note that 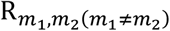 is valid only in regions where 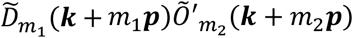 and 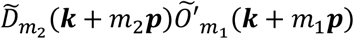 are non-zero, which are indicated by the magnitude of the produts shown in **Fig 3a** (m_1_ = 0 and m_2_ = 1). For a specific combination of (m_1_, n_1_) and (m_2_, n_2_) the residual phase, which is the phase of 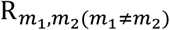, can be expressed as

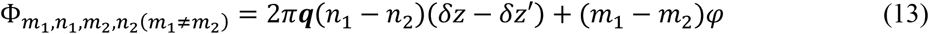

Eqn (13) is the sum of two terms. The first term is determined by the axial offset between the excitation and detection foci, will be zero if δ*z* = δ*z′* (**Fig 3b and 3c**), and is a function of *k*_*z*_ (because it contains the *k*_*z*_-oriented vector ***q***. The second term is due to the lateral phase *φ* in the illumination pattern and is a constant across the overlap region (the same as eqn (10)). This has two important implications for the reconstruction process. The first is that it creates a need to explicitly account for the axial phase offset during reconstruction. For light sheet illumination, the delta functions of the hexagonal lattice in the pupil are extended into lines or Gaussian profiles along the kz direction, corresponding to axial confinement of the light sheet at the sample. This complicates the simplified version of residual phase shown in eqn (13) and makes it challenging to derive an analytical solution. In practice, to extract the axial offset δ*z* from the sample images, we can collect a gallery of transfer functions with different known axial offsets δ*z′*, and then identify for which measured offset 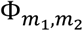 reaches a minimum value at δ*z′* = δ*z* (**Fig 3d**). The corresponding transfer functions can then be used to reconstruct the SIM images and will yield minimized artifacts due to axial misalignment between the excitation and detection focus. The second implication of eqn (13) is that it shows that the conventional method for estimating the lateral phase offset *φ* as described in eqn (10) via complex linear regression will fail as the residual phase 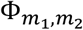 is no longer constant in the overlap region as assumed by conventional SIM reconstruction. As illustrated in **Fig 4**, the residual phase Φ is constant only when the first term in Φ is zero (δ*z* = δ*z′*, **Fig 4a**). Whenever δ*z* ≠ δ*z′*, Φ will depend on ***q***(*n*_1_ − *n*_2_). This means that the ratio 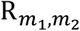 is no longer a constant and 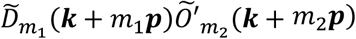 is no longer linearly related to 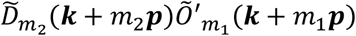. Therefore, complex linear regression applied in conventional SIM reconstruction will fail. In the following sections, we address both of these aspects, by generating a 2D map of residual phase 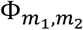 that is composed of δ*z* and *φ* and use the minimum to simultaneously extract the axial and the lateral offset needed for proper reconstruction (**Fig 5a**).

**Figure 4:**
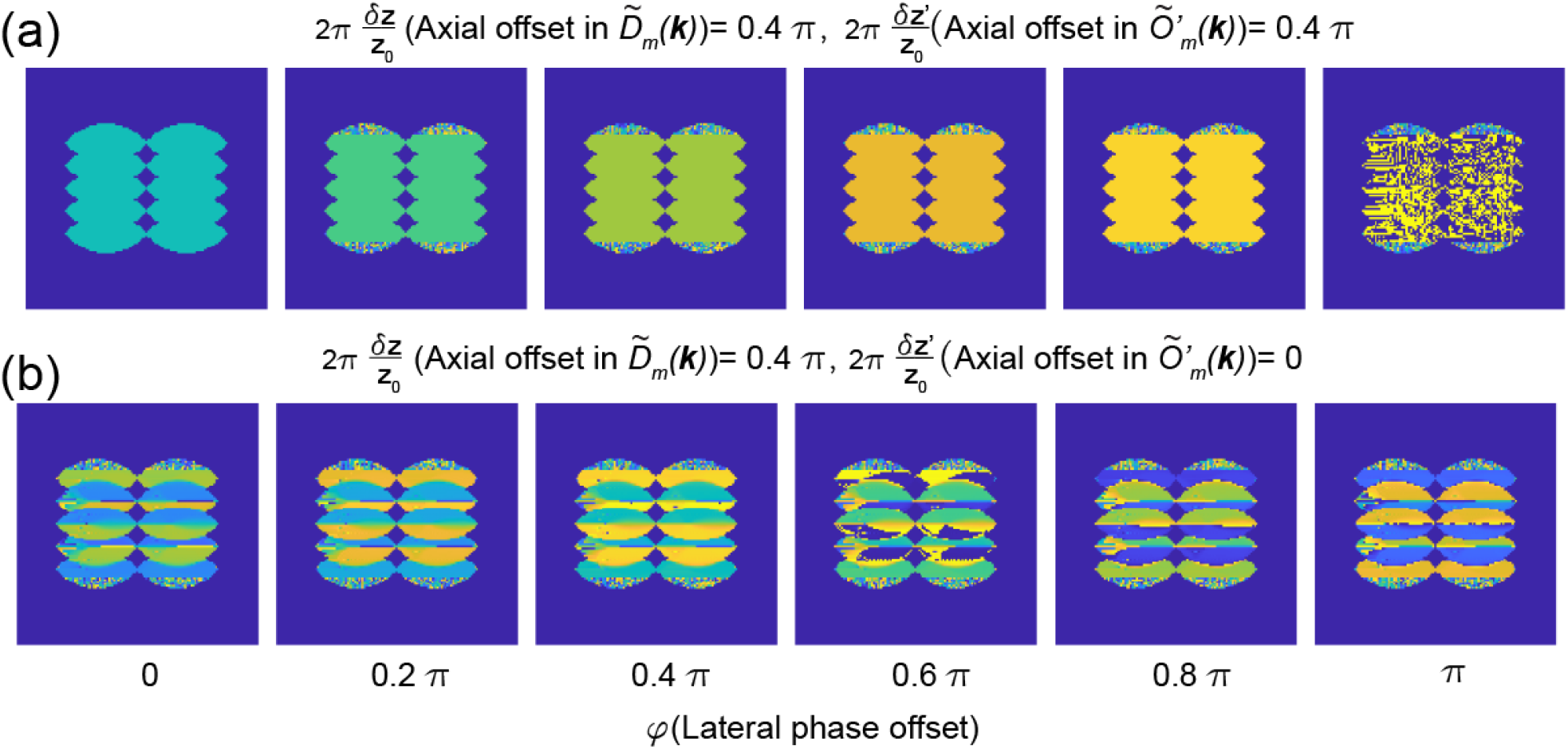
Mixing between axial and lateral phase terms in the overlap regions. **(a)** Residual phase computed via eqn (12) when both the SIM image 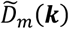 and simulated transfer functions *Õ′*_*m*_ (***k***), have a 0.4 π axial offset between the excitation and detection objectives. Different columns show the effects of different lateral offsets in the excitation pattern *Ĩ*(***k***) used when simulating the data 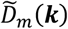 and transfer functions *Õ′*_*m*_(***k***). **(b)** Same as **(a)** but when utilizing a simulated transfer function with no axial offset.

**Figure 5:**
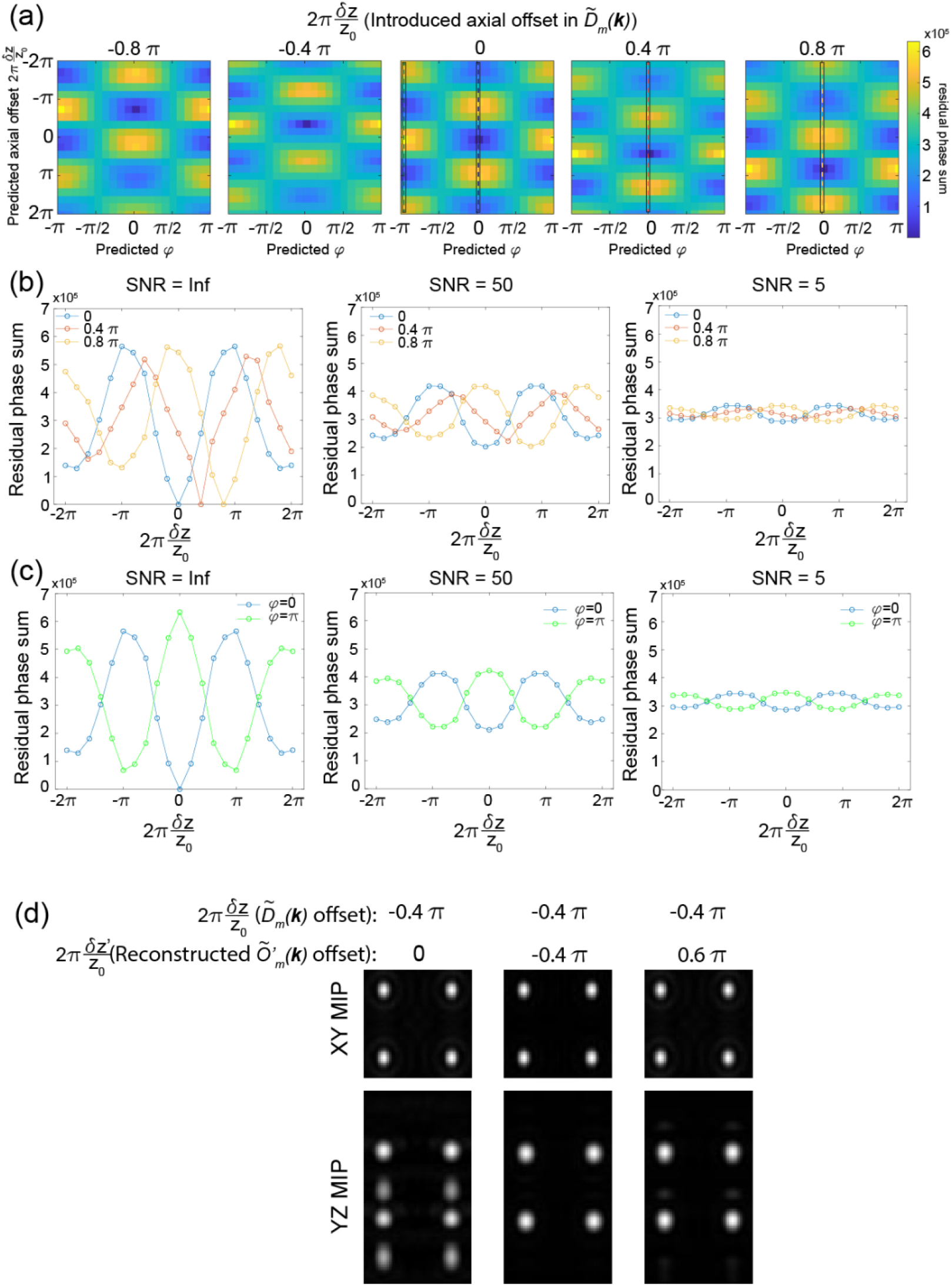
Retrieving the axial offset between excitation and detection planes in simulated data. **(a)** 2D maps of the summed residual phase in the overlap regions with different simulated axial (vertical axis) and lateral (horizontal axis) offsets in the transfer functions *Õ′*_*m*_(***k***) used for reconstruction. The simulated applied axial offset in the SIM image 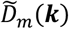 is shown on the top of each image. **(b)** Line profiles in **(a)** under different signal to noise ratios (SNR) assuming that the lateral offset between 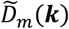 and *Õ′*_*m*_(***k***) is zero. Different colors indicate different applied axial offset in the SIM image as indicated by the dashed lines in **(a)**. Blue: no axial offset, red: 0.4 π axial offset, orange: 0.8 π axial offset. **(c)** Line profiles in (a) under different SNR when the lateral offset between 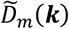 and *Õ′*_*m*_(***k***) is either zero (blue) or π (green). This illustrates the residual phase differences between the optimal pattern and the local minimum that is ½ period off both axially and laterally. **(d)** Reconstructed SIM image of four beads that were simulated with a −0.4 π axial offset between the excitation illumination and the detection focal plane. The different columns show the effects of reconstructing with transfer functions that assume no axial offset (left), the correctly identified axial offset (middle), and an axial offset that is ½ period off laterally and axially (right).

### 2.2 Simulation and image analysis

#### 2.2.1 Simulation of structural illumination microscopy images

All simulated datasets were constructed in Matlab (The Mathworks) using the previously published code base^4^. To adapt this code to generate SIM images, we generated the detection point spread function using the Debye approximation. Specifically, we first simulated the back pupil of a cone in a sphere with NA=1.0 and a refractive index of 1.33, and then Fourier transformed it into real space. We simulated the excitation point spread function following the similar approach but using a pupil of six evenly spaced vertical lines within an annular ring (**Fig 1b**, NA = 0.55/0.5). We simulated five copies of excitation PSFs whose lateral starting phase offset *φ* evenly covers the entire period by introducing a phase gradient along k_x_ in the pupil function. Because the physics of image formation are continuous, we up sampled by a factor of 2 when simulating the forward imaging process. The excitation OTF of the hexagonal pattern was dissected into different lateral orders as in eqn (3). We then followed eqn (5) and (6) to simulate the raw SIM images. Briefly speaking, the axial components of the excitation PSF *I*_*m*_(*r*_*z*_) were multiplied with the detection PSF *H*(*r*) and the lateral components of the excitation OTF *J*_*m*_(*r*_*x*_, *r*_*y*_) were multiplied with the Fourier transform of sample information *S*(***r***). The Fourier transform of *HI*_*m*_(***r***) and *J*_*m*_*S*(***r***) were then multiplied in frequency space and Fourier transformed back to real space to generate raw images *D*_*m*_(***r***). Different orders of *D*_*m*_(***r***) were then summed to form the final raw image *D*(***r***) which was downsampled to replicate image discretization onto camera pixels. To simulate different axial offsets of the excitation pattern, we applied a phase gradient along k_z_ respectively in the pupil function when we simulated the excitation PSF. Further details and a step-by-step walkthrough of the simulation process are provided in a Matlab Live script included in the accompanying Github repository.

#### 2.2.2 Simulation of randomly distributed beads

We simulated 3D volumes of randomly distributed bead images using the approach described in Shi et al^4^. In brief, we first simulated a 3D volume with points randomly distributed as *S*(***r***) in eqn (6). We then follow the pipeline above to simulate the corresponding SIM images. To simulate images of different SNR, we added a Gaussian noise floor in the simulated images.

#### 2.2.3 SIM reconstruction

SIM reconstruction was performed using code from cudasirecon^15^, following algorithm described in Gustafsson et al^7^ and using the parameters provided in the settings file in the accompanying Github repository.

### 2.3 Experiments

#### 2.3.1 Microscopy setup

The optical path for latticeSIM is based on a modified version of the instrument described in Chen et al^2^. Key modifications relevant to this work are the use of a 0.6 NA excitation lens 404 (Thorlabs, TL20X-MPL), and 1.0 NA detection lens (Zeiss, Objective W “Plan-Apochromat” x20/1.0, model # 421452-9800).

LatticeSIM was operated with a hexagonal lattice pattern with a maximum and minimum NA of 0.55 and 0.5 at the back pupil respectively. This pattern yield a period of 1.202 µm in x and 2.13 µm in z. For each z plane, we collected five copies of images, each with illumination pattern shift 0.24 µm in x and covering the entire 1.202 µm period.

#### 2.3.2 Immunofluorescence

We cultured retinal pigment epithelium (RPE) cells (RRID: CVCL_4388, ATCC) in Dulbecco’s Modified Eagle Medium (Gibco 11965-092) with 10% FBS (VWR: 1500-050) and 1% (v/v) 10,000 U/ml Penicillin-Streptomycin (Gibco 15140-122). For fixed cell imaging, we incubated RPE cells with either 250 nM mitotracker orange (Thermo Fisher, M7510) in culture media for mitochondria staining or a 1000x dilution manufacturer recommended stock concentration of SPY555-DNA (Cytoskeleton inc.) in culture media for histone staining for 30 min. We then fixed cells with 4% Paraformaldehyde (Electron Microscopy Sciences, 15710) and 8 nM/ml sucrose (Sigma, S7903) in cytoskeleton buffer (composed of 10 mM MES, 138 mM KCl, 3 mM MgCl and 2 mM EGTA) for 20 min at room temperature.

#### 2.3.3 Hydrogel preparation

Polyacrylamide substrates containing fluorescent beads were prepared as described in Tse and Engler et al^16^. Briefly, a mix of 40 % acrylamide and 2% Bis-Acrylamide was prepared in 10 mL diH20 to an estimated stiffness of 8 kPA. 1/1000 by volume of carboxylate red fluorosphere fluorescent beads (580 nm/605 nm excitation/emission wavelength, Thermo Fisher, F8801) were added to 495 microliters of the mix and placed in a vacuum dessicator for 15 minutes to reduce dissolved oxygen before addition of 5 uL of 10% by weight of APS (freshly prepared in diH_2_O) and 0.5 uL of TEMED. Immediately, 30 microliters of solution was carefully pipetted onto a 25mm glass coverslip that was previously activated with 97% APTES and 0.5% Glutaraldehyde and then a 12 mm glass coverslip was placed on top of the solution. After polymerization, the coverslip sandwich was immersed in diH_2_O water for five minutes before a razor blade was used to carefully peel the 12 mm glass coverslip off of the 25 mm coverslip. The PAA substrate on 25mm coverslip was then immersed in diH20 and stored in a 4° C fridge until use.

#### 2.3.4 Measurements of light sheet offset in hydrogel

To measure the extent to which a light sheet is deflected when imaging through thick hydrogels, we embedded 100 nm diameter red fluorescent beads (580 nm/605 nm excitation/emission wavelength, Thermo Fisher, F8801) inside the hydrogel. To measure the light sheet offset at different depth in the sample, we followed a similar approach as described in Shi et al^4^. Briefly speaking, we first axially scanned the excitation profile relative to the detection focal plane and plotted the integrated fluorescence signal from a small (3 pixel) region around each bead in the field of view. The peak of this plot defines the center of the excitation profile relative to the position of each bead underneath the sample. We then determined the position of each bead relative to the detection objective focal plane by scanning the coverslip together with the light sheet illumination along the optical axis of the detection objective (equivalent to widefield illumination). The offset of the excitation pattern relative to the detection objective focal plane is then computed from these plots by comparing the positions of the light sheet relative to the bead and the position of the bead position relative to the focal plane.

## 3 Results

### 3.1 Minimizing residual phase yields the transfer functions with the correct axial offset in simulation

To test whether our methodology can reliably extract the axial misalignment of the excitation pattern from simulated images, we first simulated a transfer function library *Õ′*_*m*_(***k***) and bead images 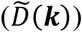 with different excitation pattern axial offsets. We quantify the sum of residual phase in the overlap region between different orders in Φ_01_ and Φ_12_ as the metric to minimize. As shown from eqn (12), the phase of Φ_01_ and Φ_12_ will be a function of axial offset δ *z′* and the lateral offset *φ*. To determine both the axial offset and lateral phase simultaneously, we search for the minimum in Φ_01_ and Φ_12_ in a 2D residual phase map corresponding to each pairing of axial and lateral offset (**Fig 5a**). As illustrated in **Fig 5b**, the axial offset corresponding to the minimum value in the 2D residual phase map matches the misalignment that we introduced in simulation. Moreover, we noticed that as the signal to noise ratio (SNR) decreases, the modulation depth of the residual phase array also decreases, making it harder to distinguish the true light sheet axial focus from the position that is one full period off (**Fig 5b**).

Another caveat of this methodology is that the aligned (blue line in **Fig 5c**) and pi offset along axial and lateral direction (green line in **Fig 5c**) are both local minimums in the 2D phase map. The difference between the two minimum also decreases as the SNR decreases (**Fig 5c, blue and green curves**). This means that with increasing noise, our method will have trouble distinguishing between transfer functions that are laterally and axially offset by half of a period rather than honing in on the correct axial offset. We next tested how this will affect the performance of SIM reconstruction. We simulated an image of 8 beads whose excitation axial focal plane is 0.4 π below the detection focal plane and we reconstructed this simulated image with the transfer functions that generated the minimum variance in residual phase (– 0.4 π offset) or the transfer functions that are a half a period off axially (0.6 π offset). As shown in **Fig 5d**, both reconstructed images show minimum artifacts compared with the image reconstructed with calibration transfer functions that did not account for axial misalignment. For axially periodic illumination patterns, there will be no difference between the two minimums as they represent different nodes in the lattice. However, the Gaussian envelope associated with the light sheet confinement differentiates these two local minimums and may cause noticeable artifacts for tightly confined beams. Therefore, in practice to minimize this effect in the experimental data, we limited the lateral offset search range for minimum between −π/2 to π/2.

### 3.2 Validation with fluorescent beads

After testing the approach via simulations, we next experimentally validated it with fluorescent beads. We deliberately introduced a known axial offset between the lattice light sheet and the detection objective focal plane, and tested whether the minimum position in the 2D phase map could recover the applied offset. As illustrated in **Fig 6a**, the minimum position in the 2D phase map within the −π/2 to π/2 lateral phase offset range (indicated by the red asterisks in **Fig 6a**) matches the introduced axial offset, thus demonstrating that our simulated results also transfer over to experimental datasets. In latticeSIM, the shape of the illuminated lattice will vary as the beam diverges along the propagation direction of the beam. Therefore, after validating that the method works when the sample is centered along the propagation direction of the excitation pattern, we next tested the performance when the sample is displaced along this axis (**Fig 6b-h**). We extracted the axial position of the minimum in the 2D phase map and compared the predicted axial offsets (blue circles in **Fig 6 c, e and g**) with those that were applied experimentally (red dashed lines). We found that at the propagation focal position (**Fig 6c**, orange line in **Fig 5b**) or 5 µm off from the propagation focal position (**Fig 6e**, green line in **Fig 5b**), we can successfully retrieve the axial offset within the range of 1.2 µm above or below the detection focal plane (one full lattice period along the axial direction is 2.13 µm), as the blue circles matched well with the red dashed lines. We investigated the performance of SIM reconstruction using the predicted transfer function from our methods. As shown in **Fig 6d and 6f**, the raw lattice light sheet images with a 600 nm axial offset between the focal plane of the excitation and the detection objectives have clear ghost copies (YZ MIP in the first column of **Fig 6d and 6f)**, and similar effects were observed in the SIM images that were reconstructed with “zero-offset” transfer functions (the 2^nd^ column of **Fig 6d and 6f**). The reconstructed images using transfer functions that were generated with the computationally predicted excitation offset (y values of blue circles in **Fig 6c** and **Fig 6e**, reconstructed images shown in the 3^rd^ column of **Fig 6d and 6f**) yield a less aberrated image. Moreover, images reconstructed with transfer functions that are half a period off axially and laterally from the predicted positions (green crosses in **Fig 6a and Fig 6c inset**) also yield comparable quality (**Fig 6d and f**, 4^th^ column).

**Figure 6:**
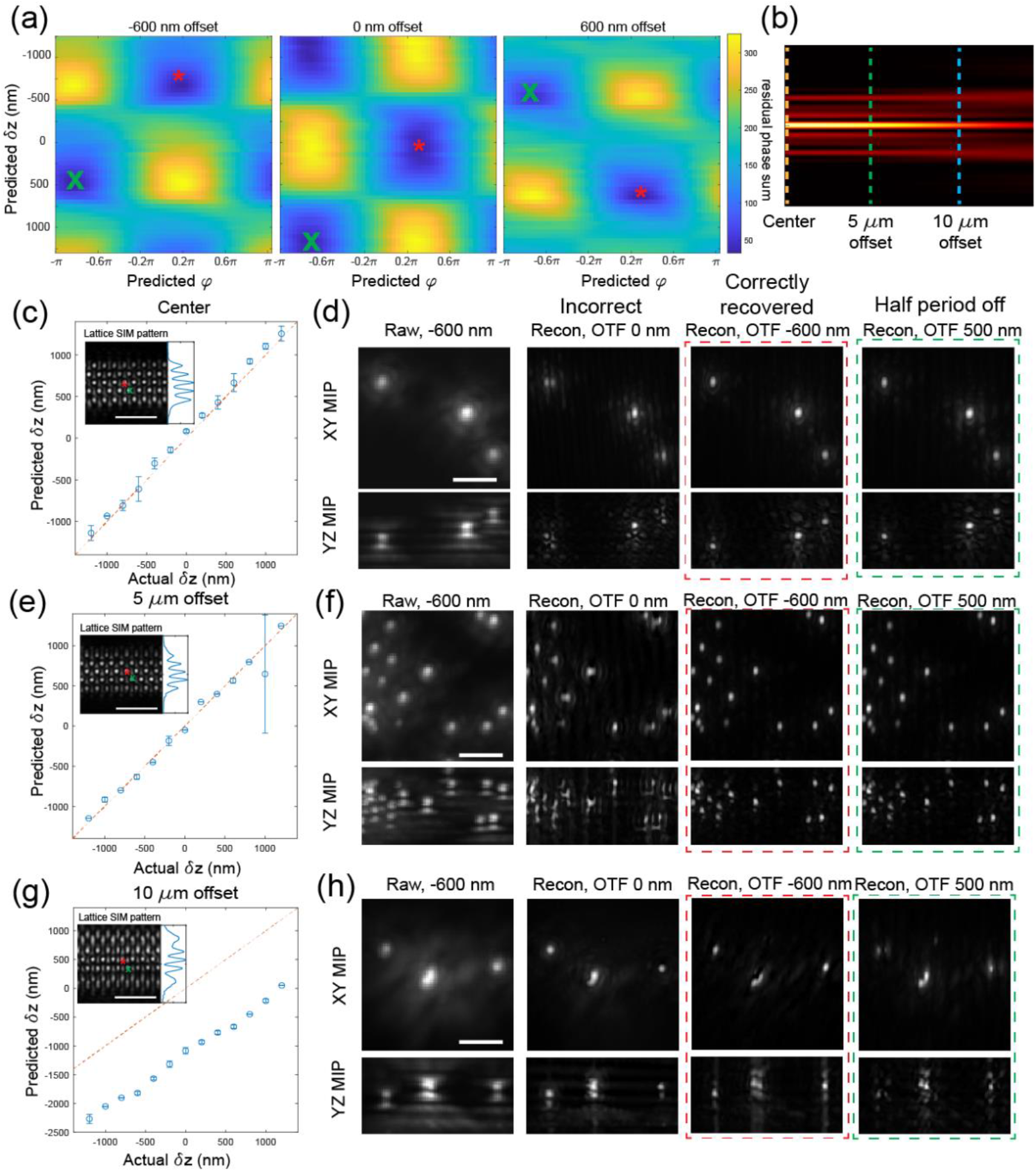
Retrieving the axial offset between excitation and detection planes in experimental data with point-like objects. **(a)** 2D maps of the summed residual phase in the overlap regions with different experimentally obtained axial (vertical axis) and lateral (horizontal axis) offsets in the transfer functions *Õ′*_*m*_(***k***) used for reconstruction. The applied axial offset in the experimentally obtained SIM data is shown on the top of each image. The red (*) show the minimum residual phase measured between −0.5 π to 0.5 π lateral offset. The green (x) shows the local minima that is located ½ period off axially and laterally. **(b)** Different offsets in the excitation pattern along the beam propagation direction. Orange: no offset, green: 5 µm offset, blue: 10 µm offset. **(c)** Comparison between the applied (horizontal axis) and predicted (vertical axis) axial offset between the excitation and detection focal planes in SIM image taken with the sample located at the center of the beam (orange line in **(b)**). Inset shows the illumination pattern *I*(***r***). Red (*) and green (x) correspond to the two minima in **(a)**. Scale bar = 5 µm. **(d)** Raw and reconstructed SIM images of fluorescent beads taken under the same settings as in **(c)** and with a −600 nm axial offset between the excitation and detection focal planes. The 1^st^ column shows the raw image, the 2^nd^ column shows the image reconstructed with transfer functions using the correctly predicted axial offset (−600 nm), the 3^rd^ column shows the image reconstructed with transfer functions whose excitation and detection focal planes are aligned, the 4^th^ column show the image reconstructed with transfer functions that utilize an axial offset that is ½ a period off from the predicted value (i.e. 500 nm vs. −600 nm). Scale bar = 2 µm. **(e, f)** same as **(c)** and **(d)** but images are taken with 5 µm offset from the beam center along the propagation direction. **(g, h)** same as **(c)** and **(d)** but images are taken with 10 µm offset from the beam center along the propagation direction.

However, we also noticed that when imaging with an excitation pattern that is 10 microns away from the focus along the beam propagation direction (**Fig 6g and h**, blue line in **Fig 6b**), the predicted axial offset locked in on one that is a half period off laterally and axially (**Fig 6g**). We suspect that this is because the aberrations in the excitation pattern for this condition actually represent a combination of a linear phase ramp due to the axial beam offset and a defocus phase term due to the propagation offset. A failure to account for the defocus phase term likely reduced the ability to distinguish between the two local minima that are one-half period away from each other. Further, as in **Fig 6h**, images reconstructed with transfer functions having the correct axial offset (3^rd^ column in **Fig 5h**) or half a period off (4^th^ column in **Fig 5h**) both yield symmetric side lobes in the YZ max intensity projection (MIP). However, we also noticed that, even at the corrected axial focus, these reconstructed images were more aberrated compared with the ones in **Fig 6d** and **f** where the sample was at the propagation focus of the light sheet. We believe that this is likely because the structured illumination pattern is more distorted compared with the ones at the focal position or 5 µm off from the focal position. Therefore the weighting among different frequency components of *Õ*_*m*_(***k***) is different from *Õ′*_*m*_(***k***), which will yield artifacts during the Wiener deconvolution step in image reconstruction. However, the fact that the side lobes are symmetric in **Fig 6h** still indicated that our method can still successfully predict the amount of axial offset given the raw image.

### 3.3 Validation with adherent cells

In order to benchmark the performance of our method on biological samples, we imaged two subcellular structures in RPE cells: mitochondria and nuclei. As illustrated in **Fig 7a and b**, our method can successfully retrieve the introduced axial excitation offset in these two structures over the range of −1.2 µm to +1.2 µm. Similar to our benchmark with beads, raw lattice light sheet images taken with a 600 nm offset between the excitation focal plane and the detection focal plane show ghost copies in the YZ MIP, and the same ghost copies appeared in the reconstructed images that were processed with “zero-offset” transfer functions (**Fig 7c and d**, 1^st^ and 2^nd^ columns). Moreover, using transfer functions with the correctly predicted axial offset again yielded an image with minimal aberration (**Fig 7c and d**, 3^rd^ column). The same trends can be observed in linecuts of the image power spectrums where the SIM image reconstructed with the corresponding transfer functions (with or without axial offset between the excitation and detection focal planes, red and blue curves in **Fig 7e**) have better high-frequency support compared to SIM images that were reconstructed with incorrect transfer functions. Overall, these results highlight that our method can successfully retrieve the axial offset from the raw latticeSIM images and yield reconstructions with minimal aberrations even when imaging complex 3D structures in cells.

**Figure 7.**
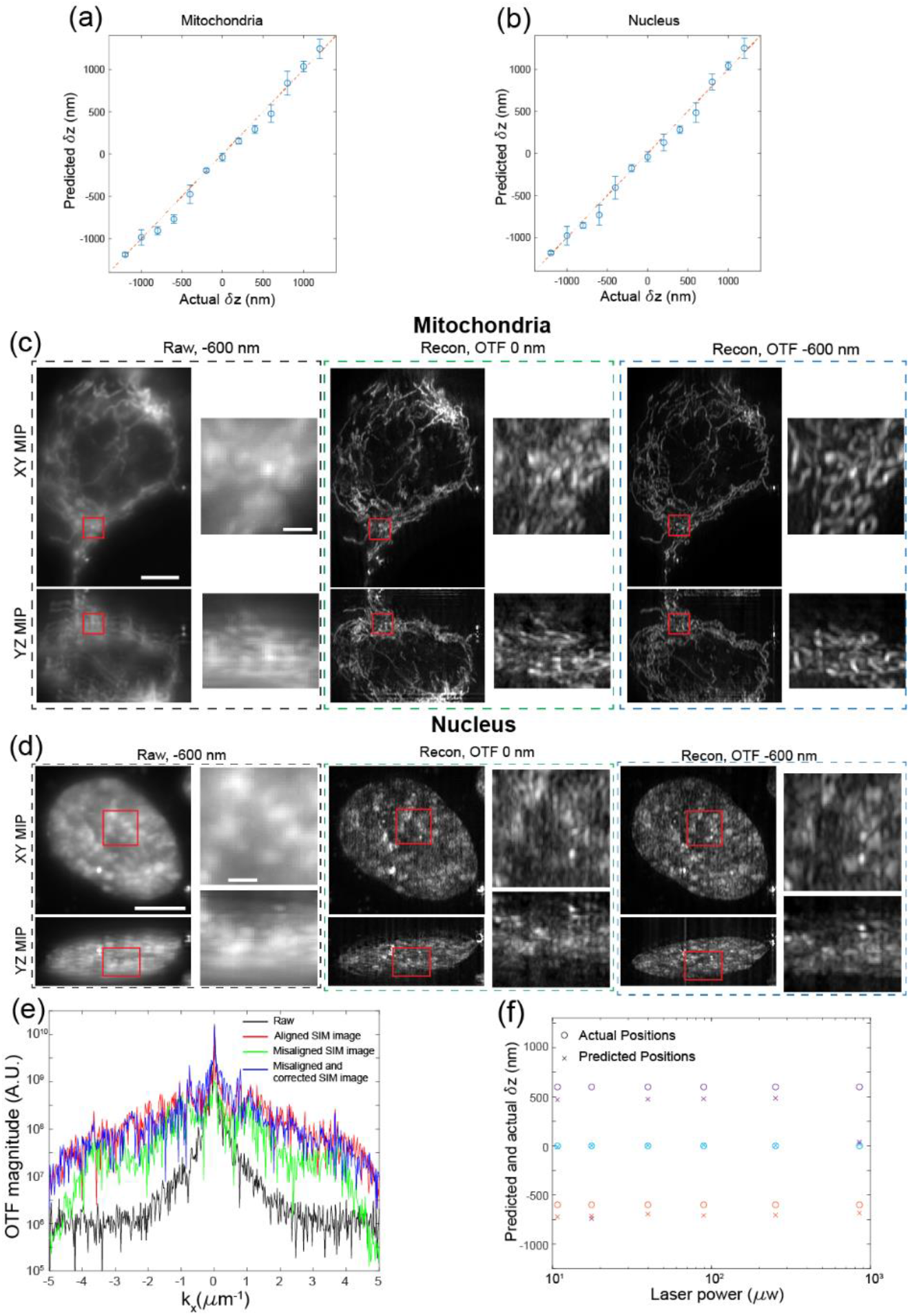
Demonstration of method with adherent cells. **(a, b)** Comparisons between the experimentally applied (horizontal axis) and computationally predicted (vertical axis) axial offset between the excitation and detection focal plane when imaging cellular mitochondria **(a)** and the cell nucleus **(b). (c)** Raw and reconstructed SIM images of mitochondria taken with the cell aligned at beam propagation focus (orange line in **Figure 6b**)) and with a −600 nm axial offset between the excitation and detection focal planes. The 1^st^ column shows the effective dithered lattice image by summing the images from each of the 5 lateral phase steps, the 2^nd^ column shows the image reconstructed with transfer functions whose excitation and detection focal planes are aligned, and the 3^rd^ column shows the image reconstructed with transfer functions that used a computationally identified axial offset of −600 nm. Scale bar = 5 µm. Insets show a zoomed in view of the red-box, inset scale bar = 2 µm. **(d)** Same as **(c)**, but with images of the cell nucleus. **(e)** Linecuts of the power spectrums for the 5-phase summed raw images (black), and images that were acquired and reconstructed under the following settings: 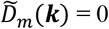 nm and *Õ′*_*m*_ (***k***) = 0 nm (red), 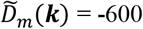 nm and *Õ′*_*m*_ (***k***) = 0 nm (green), 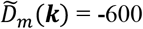 nm and *Õ′*_*m*_ (***k***) = −600 nm (blue). **(f)** Comparisons between the applied (circles) and predicted (crosses) axial offsets between the excitation and detection focal plane in images of mitochondria taken with different laser power at the back pupil of the excitation objective. The test was run at three different experimentally applied offsets: −600 nm (orange), 0 nm (cyan), and +600 nm (purple).

Another factor that will affect the performance of our method is SNR. Because SNR may be low when imaging biological samples where phototoxicity and photobleaching need to be minimized, we tested our methods with cellular mitochondria using different laser powers. As shown in **Fig 7f**, the accuracy of our approach did not degrade substantially as the applied laser power decreased. This indicated that our method is robust across SNR, at least over the range measured between 4 to 25.

### 3.4 Validation with fluorescent beads in 3D hydrogel

Thus far, we’ve validated that our method can posteriorly extract the axial offset of the excitation pattern from raw SIM images of beads and adherent cells. However, these structure are still relatively thin. To address how our algorithm would perform in thicker samples, we next tested its performance using fluorescent beads embedded within thick 3D hydrogels. Because beads in the imaged hydrogels span multiple locations along the light sheet propagation direction and to exclude possible aberrations from this aspect (covered in **Fig 6)**, we only include beads illuminated by ±5µm within the focus along the light sheet propagation direction when computing the axial offset of the beam. To quantitatively validate whether the prediction from our methodology is correct and assess additional artifacts from imaging through the hydrogel, we also independently measured the offset between the excitation focus (determined by the axial profile of the lattice light sheet) and the detection focus (determined by the axial profile of the bead illuminated by widefield illumination). As shown in **Fig 8a**, our method can correctly extract the sample induced excitation axial offset within 5 µm below the hydrogel surface. However, our method tends to overestimate this offset for beads more than 10 µm below the hydrogel surface. This mismatch is likely attributed to the distortion of the illumination pattern (and possibly the detection PSF) caused by the hydrogel refractive index mismatch with the media (**Fig 8b**). The excitation aberration becomes visually apparent when the depth is larger than 10 µm. Therefore, the assumption of our method that, other than axial or lateral translations, the same excitation pattern and detection PSFs are applied to take the both the transfer function library *Õ′*_*m*_(***k***) and the sample images 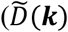 is no longer valid.

**Figure 8.**
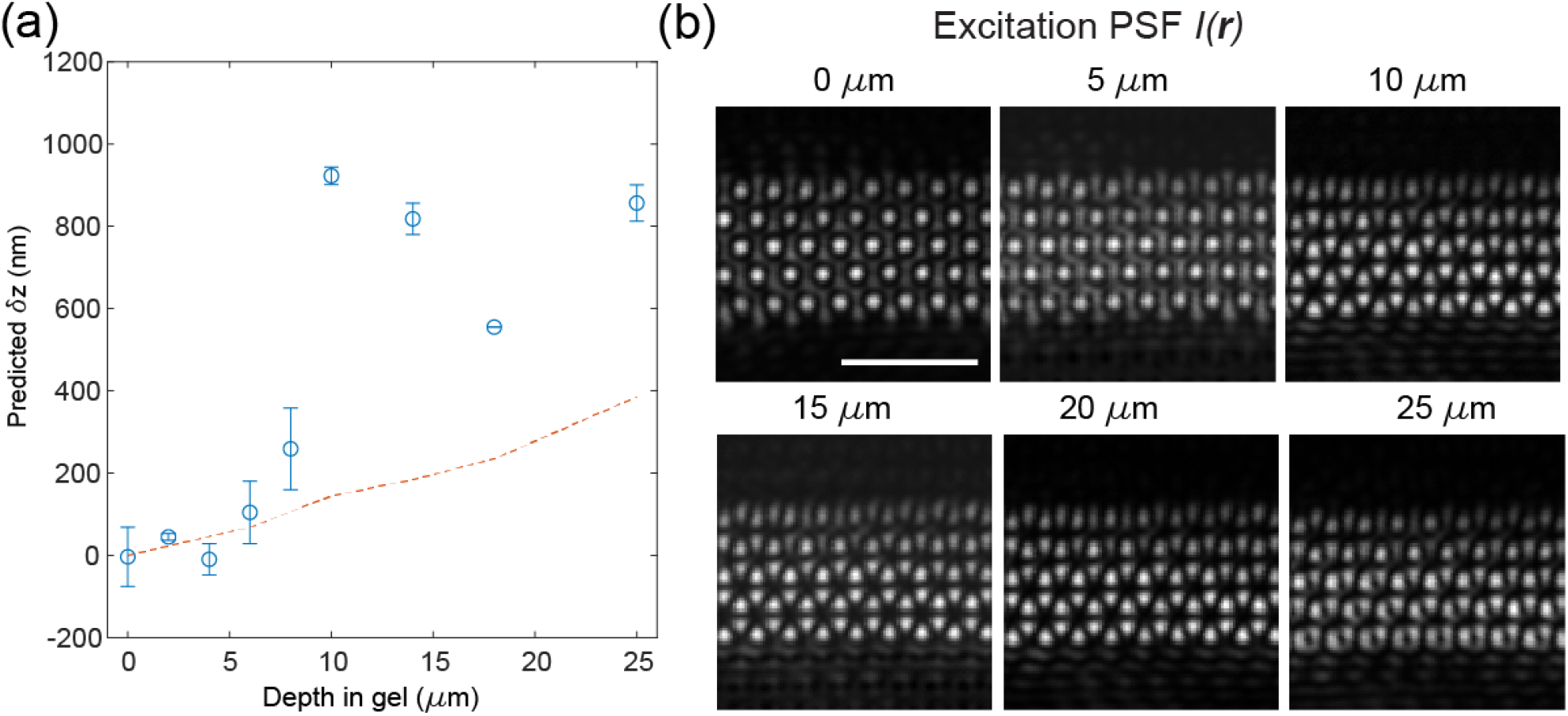
Demonstration of method in 3D hydrogels. **(a)** Comparison between the experimentally measured (red dashed lines) and computationally predicted (blue circles) axial offset between the excitation and detection focal plane in images of fluorescent beads taken at different depths in a 3D polyacrylamide hydrogel. In this case, no offset was experimentally applied, but the hydrogel introduced a shift in the excitation pattern that varies as a function of depth. **(b)** Experimentally measured excitation pattern (*I*(***r***)) taken at different depths in the hydrogel. Scale bar = 5 µm.

## 4 Discussion

In this paper, we introduce a posterior method for extracting the axial offset between the excitation light sheet and the detection focal plane in latticeSIM. We validated this method through simulations and experiments, employing fluorescent beads, adherent cells, and 3D hydrogels. Our demonstrations illustrate that identifying the transfer functions with the same retrieved axial offset as that of the raw SIM image minimizes artifacts in the reconstructed images. Overall, this approach enables posterior correction of system or sample-induced axial offsets in latticeSIM. Importantly, it only necessitates a gallery of transfer functions with varying axial offsets that can be acquired experimentally or computationally generated, without the need for additional optical components in the microscope.

One caveat of our method is that with decreasing SNR, it can become challenging to differentiate between the axial focal position of the light sheet and the position that is half a period off (see **Fig 5a** and **Fig 6a**). This challenge is particularly pronounced for hexagonal lattice patterns with a small difference between the minimum and maximum NA of the bounding pupil mask. The x-averaged axial intensity profile shown in **Fig 1c** and **Fig 6c** inset reveals that the center peak (representing the true axial focal position) and the two side lobes (representing axial positions that are half a period off) possess similar magnitudes. Consequently, this explains why images reconstructed with the correct axial offset or those shifted by half a period exhibit comparable image qualities. We expect that lattice patterns with more pronounced differences in intensity between the center and side lobes (such as a square lattice or a hexagonal lattice with a larger N.A. gap) will exhibit greater disparities between the two minima corresponding to the center peak and the side lobes, making them easier to distinguish via our approach, but also more sensitive to artifacts when it fails and inaccurately identifies a half-period shifted pattern.

Another limitation of our approach is that it assumes that the same illumination pattern is applied in both the SIM image and during calibration of the transfer function library. This necessitates that, other than translational shifts, the OTFs of the excitation pattern and detection objective remain identical between the two instances; failure to meet this assumption can result in artifacts during reconstruction. For example, alterations in the magnitude distribution within the excitation OTF relative to that of the transfer function library will lead to incorrect normalization during SIM reconstruction and introduce artifacts. Furthermore, sample-induced aberrations that cause distortion of the lattice pattern will result in erroneous predictions by our method, as demonstrated in the 3D hydrogel where aberrations induced by a refractive index mismatch distorted the hexagonal illumination pattern. Finally, even in systems with adaptive optics, correction for aberrations across the entire biological sample can be challenging. Thus, a single posterior correction may not be valid over an entire image. In these cases, we postulate that our method could be combined with tiled SIM reconstruction^17^ to account for this spatially variant offset.

Nevertheless, we demonstrate here that the method performs well for adherent cells where the light sheet offset can be considered to be constant across the usable field of view. As these examples have made up the majority of use cases for latticeSIM, we envision that our method will be a valuable addition to restore image quality even under non-ideal experimental conditions. Furthermore, if we view the axial offset of the excitation beam as a phase aberration (e.g. Noll Zernike indices 2 and 3), then we envision that extensions of this method may be used to correct for higher-order or even arbitrary phase aberrations in the excitation beam as long as they can be represented in a gallery of experimentally obtained or simulated transfer functions and used for residual phase minimization. This advancement would further enable tiled reconstruction for latticeSIM experiments in complex 3D specimens.

## Disclosure

W.R.L. is an author on patents related to Lattice Light Sheet Microscopy and its applications including:

U.S. Patent #’s: US 11,221,476 B2, and US 10,795,144 B2 issued to W.R.L. and coauthors and assigned to Howard Hughes Medical Institute. Y.S. and T.A.D. declare no competing interests.

## Code and Data Availability

Due to the inordinate size of the image data (∼700GB), it is not currently feasible to deposit this into a central repository; however, all datasets underlying the results in this manuscript are available from the corresponding author upon request. To the extent possible, the authors will try to meet all requests for data sharing within 2 weeks from the original request.

Code to reproduce the results shown in this manuscript, a small demonstration dataset, and the setting file for SIM reconstruction are available at: https://github.com/legantlab/SIM_Aligment

## Acknowledgements

We thank Eric Betzig for helpful discussion. This work was funded in part by grants from the National Institutes of Health (1DP2GM136653) awarded to W.R.L.. W.R.L. acknowledges additional support from the Searle Scholars program, the Beckman Young Investigator Program, and the Packard Fellowship for Science and Engineering.

